# macrosyntR : Drawing automatically ordered Oxford Grids from standard genomic files in R

**DOI:** 10.1101/2023.01.26.525673

**Authors:** Sami El Hilali, Richard R. Copley

## Abstract

Macrosynteny refers to the conservation of chromosomal to sub-chromosomal domains across species and its conservation can provide insight on the evolution of animal genomes. Pairwise comparison of de-novo assembled genomes based on predicted protein sequences often use a graphical visualization called an Oxford grid. We implemented an R package to draw Oxford grids from standard genomic file formats. The package can automatically order the chromosomes, to improve interpretability, and is thus helpful for both exploratory data analysis and production of publication quality graphics.

## 1. Introduction

Rearrangements of ancestral metazoan chromosomes have shaped the karyotypes of current animals. Studies based on distantly related animals have suggested good conservation of macrosynteny, that is, conservation of gene content within chromosomal or sub-chromosomal domains, but not necessarily conservation of precise gene order (Simakov *et al*., 2022). Synteny analyses are becoming increasingly common in comparative genomics studies of eukaryotes (Tang *et al*., 2008; Wang *et al*., 2017).

In the context of efforts like the Earth BioGenome Project (Lewin *et al*., 2018) newly released genome assemblies of non-model organisms are reaching chromosomal scales. These data will provide a valuable resource to build a comprehensive framework of the evolution of metazoan chromosomes. At the same time many computational tools (reviewed in (Lallemand *et al*., 2020)) have been implemented to allow for the analysis of the macro and microsynteny conservation.

In Metazoa, the pairwise comparison syntenic relationships often uses a graphical visualization called an Oxford Grid, a two-dimensional dot plot showing genomic loci of two species, each being assigned an axis, and every dot representing a correspondence in DNA content (Edwards, 1991). More recently, this concept has been used to show the chromosomal correspondences of inferred gene ortholog pairs (Wang *et al*., 2017; Simakov *et al*., 2020; Martín-Durán *et al*., 2020; Brasó-Vives *et al*., 2022).

The Oxford Grid is a straightforward visualization tool to compare the genome-wide distribution of orthologous genes in high quality genomes. We have implemented macrosyntR, an open source R package to automatically identify, order and plot the relative spatial arrangement of orthologous genes on Oxford Grids. It features an option to use a network-based greedy algorithm to cluster the sequences that are likely to originate from the same ancestral chromosome.

## 2. Description

macrosyntR is an R package that features 5 functions covering all actions needed to go from input files to the drawing of Oxford Grids. A stable version is available on CRAN (https://CRAN.R-project.org/package=macrosyntR) and the developmental version is hosted on github at: https://github.com/SamiLhll/macrosyntR

### 2.1. Input DATA

The data required are a table of orthologous genes, that is a two column table containing sequence names of pairs of orthologous genes in the two species, and one BED file (https://www.ensembl.org/info/website/upload/bed.html) per species to specify gene order, each composed of the three mandatory fields Chrom, ChromStart, ChromEnd and the additional field seqName. The sequence names of the two files must match with sequence names of the first and second column of the table of orthologs.

BED files can be easily generated from other common formats such as GTF and GFF available on public databases or output by gene prediction software.

### 2.2. Computing the significantly conserved macrosynteny blocks

The function *‘compute_macrosynteny()’* builds a contingency table of the number of shared orthologs in each possible chromosome pairing. The significant associations are computed using Fisher’s exact test as described in (Simakov *et al*., 2020). The result is a data.frame object containing the number of orthologs, multiple-testing adjusted p-values that can be visualized using the function *‘plot_macrosynteny()’* (Figure 1A).

**Figure 1 :**
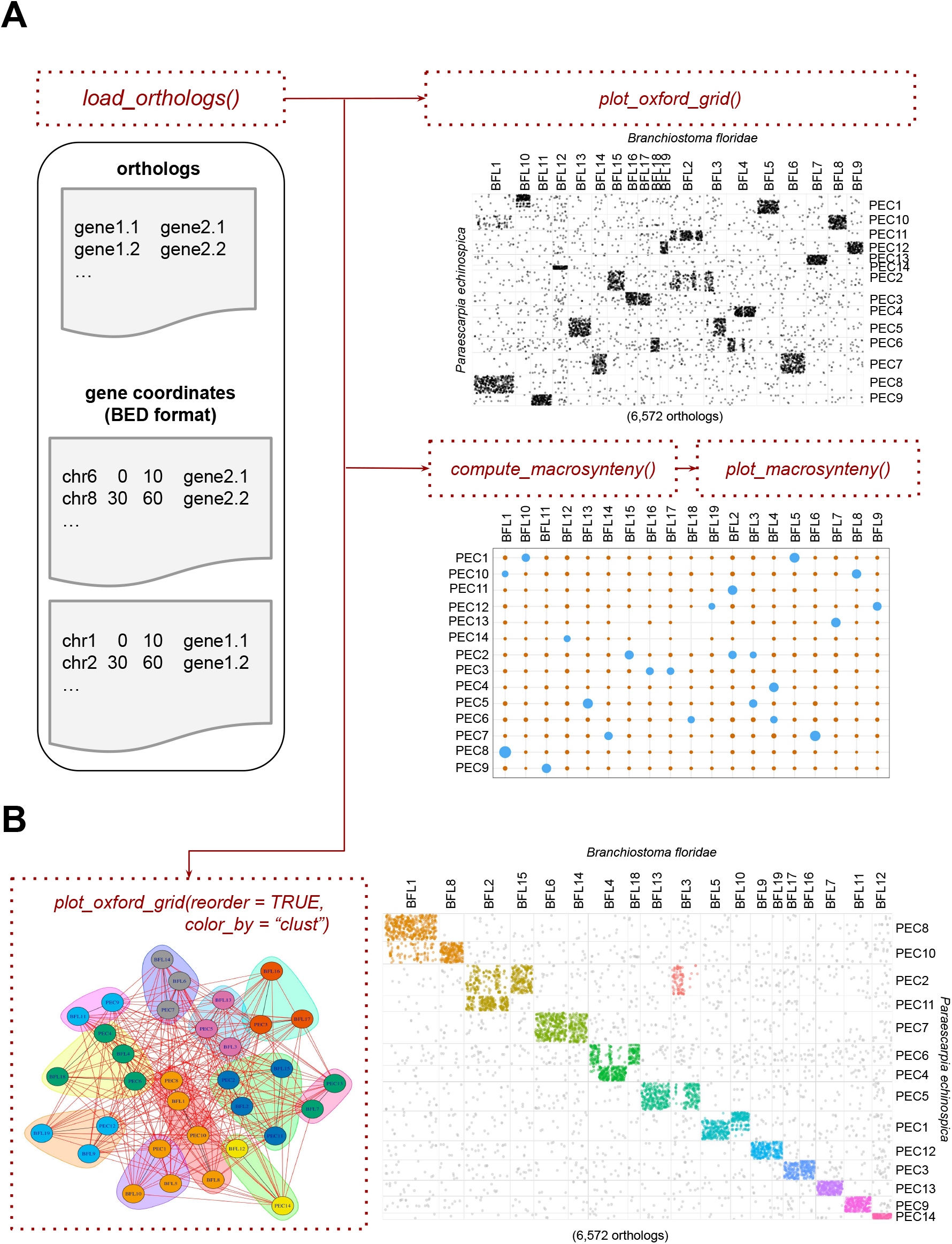
A. macrosyntR workflow for the drawing of an oxford Grid and a macrosynteny plot. B. Detection of communities to order the chromosomes. Each vertice corresponds to a chromosome from one of the two species. The edges are weighted by the amount of shared orthologs. A fast greedy algorithm is applied to detect the communities before plotting a reordered Oxford Grid.

### 2.3. Visualizing the orthologs on an Oxford grid

The Oxford grid is drawn using the function *‘plot_oxford_grid()’*. Each dot corresponds to an orthologous pair (from the function *‘load_orthologs()’*) with x and y coordinates being the relative order index on the chromosomes taken from the BED files. The dots are organized in squared facets, corresponding to the chromosomes (entry Chrom from the BED files). The limits of these squares are proportional to the total amount of genes on each scale. Many features are customizable, such as the labels, the color, size and transparency of the dots.

### 2.4. Clustering the conserved macrosynteny blocks

It is helpful to reorder the facets to interpret the results especially when analyzing fragmented genomes. Network-based methods such as hierarchical clustering have been proposed in previous studies (Putnam *et al*., 2007; Martín-Durán *et al*., 2020).

We implemented a clustering algorithm that allows for a good interpretability of the results in a limited amount of computational time (Figure 1B). Data are modeled in a network where every chromosome from the two species is modeled as a node, and weighted edges are created between every pair of nodes. The weights correspond to the number of shared orthologs between the two nodes that they connect. The linkage groups are retrieved by organizing the chromosomes into communities using the greedy algorithm implemented in the function *‘cluster_fast_greedy()’* from igraph v.1.3.0 (Csardi *et al*., 2006) It finds the local optimal communities at each step and is designed to handle large networks with an approximate linear complexity (Clauset *et al*., 2004).

The ordering is then performed by decreasing size of both the “communities”, and the chromosomes (in amount of orthologs) from left to right (species 1) and top to bottom (species 2).

This algorithm performs well even when comparing a chromosomal level genome assembly with a more fragmented one. Computation time can be reduced by using the argument ‘keep_only_significant’.

### 2.5. Example using public datasets

To demonstrate the use of the package, we compare two publicly available datasets. We selected the gene predictions of the lancelet *Branchiostoma floridae* (Simakov *et al*., 2020) and the deep-sea tubeworm *Paraescarpia echinospica* (Sun *et al*., 2021). Orthologs were calculated as reciprocal best hits using Diamond Blast (Buchfink *et al*., 2021) called through rbhXpress (El Hilali, 2022). Two R command lines are then enough to draw an Oxford Grid (Figure 1A). The default order is the chromosomal alphabetical order. We can improve the interpretability by using the clustering method (Figure 1B) and track the chromosomal fusions that have shaped the karyotypes of these species.

## Availability

A stable version is available on CRAN (https://CRAN.R-project.org/package=macrosyntR) (“*install.packages(‘macrosyntR’)*”) and the developmental version is hosted on github at: https://github.com/SamiLhll/macrosyntR

## Acknowledgements

We thank Mohamed Khamla (LBDV) for drawing of the Hex Logo.

